# EC-isHCR: a rapid method for *in situ* hybridization chain reaction in diverse animal samples

**DOI:** 10.64898/2025.12.27.696653

**Authors:** Yasuhiro Kozono, Kyohei Mikami, Shunta Yorimoto, Taisei Hayashi, Hibiki Okamura, Qingyin Qian, Ryo Hoshino, Takumi Kamiyama, Yuya Sanaki, Miho Asaoka, Sonoko Ohsawa, Shota Azuma, Yuka W. Iwasaki, Makoto Hayashi, Yu Hayashi, Shuji Shigenobu, Ryusuke Niwa, Satoru Kobayashi

## Abstract

The *in situ* hybridization chain reaction (isHCR) visualizes RNA across multiple spatial scales, from organs to subcellular structures, in diverse samples. We previously proposed a rapid protocol, EC-isHCR, for *Drosophila* embryos and ovaries. Whether EC-isHCR retains the features of conventional isHCR, including wide-spatial-scale analyses in various samples, however, has remained unclear. Here, we show that EC-isHCR enables robust RNA detection in a broad range of samples, such as whole-mount fruit fly, parasitoid wasp, and aphid preparations; paraffin sections of trout; frozen mouse sections; and human cultured cells. Moreover, EC-isHCR enabled detection of subcellular RNA localization. EC-isHCR also visualized co-localization of RNA with phase-separated condensates in fruit fly embryos and detected the protrusion-enriched mRNA in HeLa cells. To broaden the applicability of EC-isHCR, we developed an automated probe design tool (https://github.com/ShuntaYorimoto/hcrkit). By combining this tool with EC-isHCR, we provide a fast and versatile framework to visualize mRNAs. This framework will help reduce the barrier to using fast isHCR and thereby facilitate research across diverse areas of the life sciences.

**Graphical abstract:** 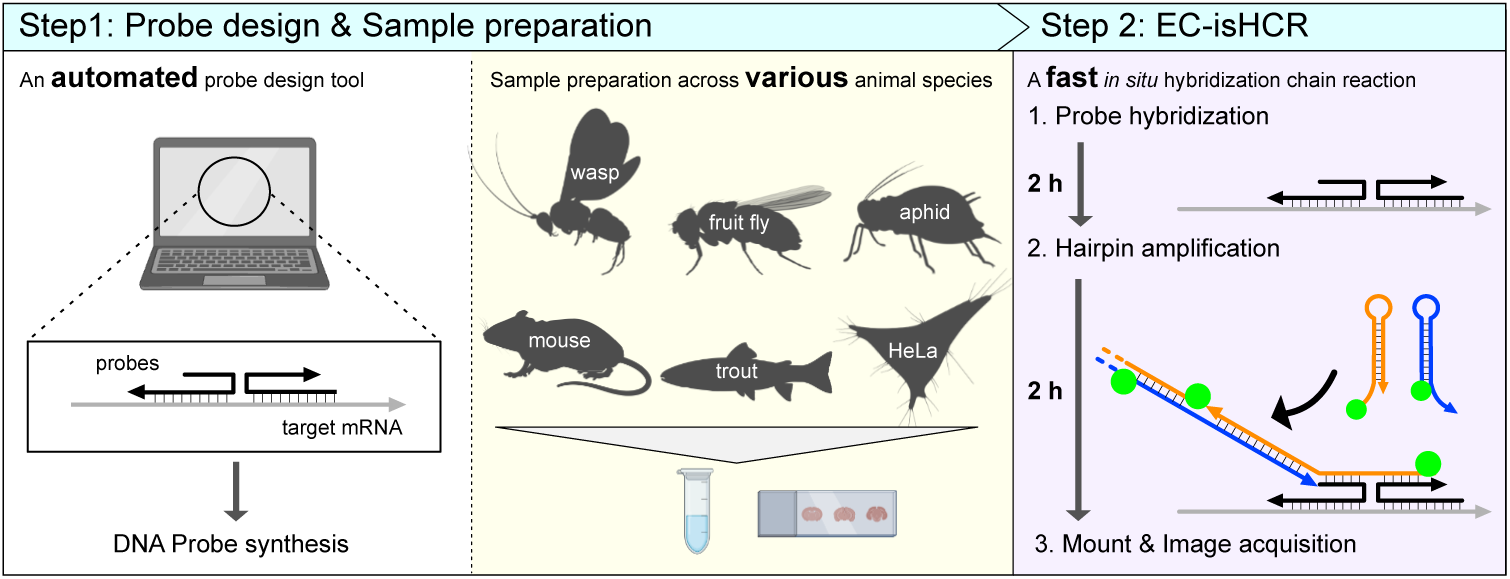

**Highlights:**

- EC-isHCR enables rapid acquisition of high-contrast images.
- EC-isHCR preserves features of conventional isHCR, including versatile sample compatibility and high-resolution imaging.
- An automated probe design tool was developed for EC-isHCR.
- EC-isHCR/probe tool framework will help reduce the barrier to using fast isHCR.

## 1. Introduction

The *in situ* hybridization chain reaction (isHCR) is an improved *in situ* hybridization (ISH) method that enables RNA detection across multiple spatial scales, from organs to subcellular structures [1–3]. By preserving spatial information, isHCR reveals differences in RNA expression among organs and cell types, providing critical insights into gene regulation and functions of specific organs and cell types. Visualizing the subcellular localization of RNAs allows assessment of whether RNAs are distributed throughout the cell or condensed with other RNAs or proteins in specific subcellular regions, providing insight into RNA dynamics within cells. isHCR offers several features, such as high contrast and compatibility with a wide range of sample types [1,2]. Thus, isHCR is an effective method for validating transcriptome data at the spatial scale of the analyzed samples [4].

Several modifications have been developed to enhance the utility of isHCR. The most broadly used approach is based on third-generation isHCR, which employs split probes [2]. Split probes hybridize to the target RNA, forming a local basis, called an initiator. The initiator anneals with fluorescently labeled hairpin DNAs, thereby tethering the target RNA to numerous fluorescent molecules through the hybridization chain reaction. Modified isHCR (a modified version of third-generation isHCR) uses shorter fluorescently-labeled hairpin DNAs to reduce reagent cost and this modification may enhance the penetration of hairpin DNAs into samples [5]. Although third-generation isHCR and modified isHCR provide powerful imaging capabilities, both methods require 3 days to stain whole-mount samples [2] and 2 days to stain sectioned samples [5], which is much longer than other ISH methods, including RNAscope and single-molecule RNA fluorescence *in situ* hybridization (smFISH) [6,7].

We previously developed ethylene carbonate-isHCR (EC-isHCR), a faster isHCR protocol by employing ethylene carbonate (EC) in combination with shorter hairpin DNA [8]. EC is a nucleic acid denaturant that efficiently promotes the hybridization reaction and reduces the hybridization time compared to formamide [9]. EC-isHCR reduces staining time to a single day, yielding high-contrast multiplex images in whole-mount samples. To date, however, EC-isHCR has only been applied to a few *Drosophila* samples with a limited spatial scale [8,10], and it has remained unclear whether EC-isHCR retains the features of conventional isHCR, including wide-spatial-scale analyses in a wide range of samples.

Here, we evaluated the versatility of EC-isHCR across various sample types and assessed the subcellular resolution of images obtained using this method. EC-isHCR demonstrated robust RNA detection across samples prepared using representative fixation methods, including whole-mount *Drosophila*, parasitoid wasp, and aphid preparations; paraffin sections of trout; frozen mouse sections; and human cultured cells. EC-isHCR also detected co-localization of mRNA with RiboNucleoProtein (RNP) granules, a subcellular structure, in *Drosophila* embryos and protrusion-enriched mRNA in HeLa cells. In addition, we developed an automated probe design tool that is applicable to a wide range of species. By combining EC-isHCR with this probe design tool, we provide a simple and fast isHCR framework for a broad range of samples and applications.

## 2. Results

### 2.1. Applicability of EC-isHCR in various whole-mount Drosophila samples

We first examined whether EC-isHCR can be applied to *D. melanogaster* larval samples. In female larval gonads, *vasa* and *traffic jam* (*tj*) mRNA are expressed in different cell types (*vasa*: germline, *tj*: intermingled cells, which are somatic stromal cells) [4]. Consistent with the previous report [8], EC-isHCR detected these mRNAs (Fig. 1A–C). We next performed EC-isHCR in adult samples. In the adult *D. melanogaster* midgut, *escargot* (*esg*) mRNA is mainly expressed in intestinal stem cells (ISCs) and enteroblasts (EBs) [11,12], and *Tachykinin* (*Tk*) mRNA is expressed in enteroendocrine cells (EECs) [13]. *esg* mRNA signals were detected in ISCs and EBs marked with *esg* reporter (*esg-GAL4>UAS-EGFP*), and *Tk* mRNA signals were observed in EECs marked with Prospero (Pros) protein (Fig. 1D–K). Additionally, we applied EC-isHCR to the male accessory gland (MAG), a reproductive organ that produces major seminal fluid components[14]. The MAG is mainly composed of main cells, which predominantly express *Sex Peptide* (*SP*) mRNA encoding a seminal fluid protein [15,16]. EC-isHCR successfully detected *SP* mRNA (Fig. 1L–O). Furthermore, we observed *SP* mRNA in the MAG of *D. sechellia*, a species phylogenetically close to *D. melanogaster* (Fig. 1P–S). Taken together with our previous results in embryos and adult ovaries [8], these findings demonstrate that EC-isHCR can visualize mRNAs in various *Drosophila* samples.

**Fig. 1.**
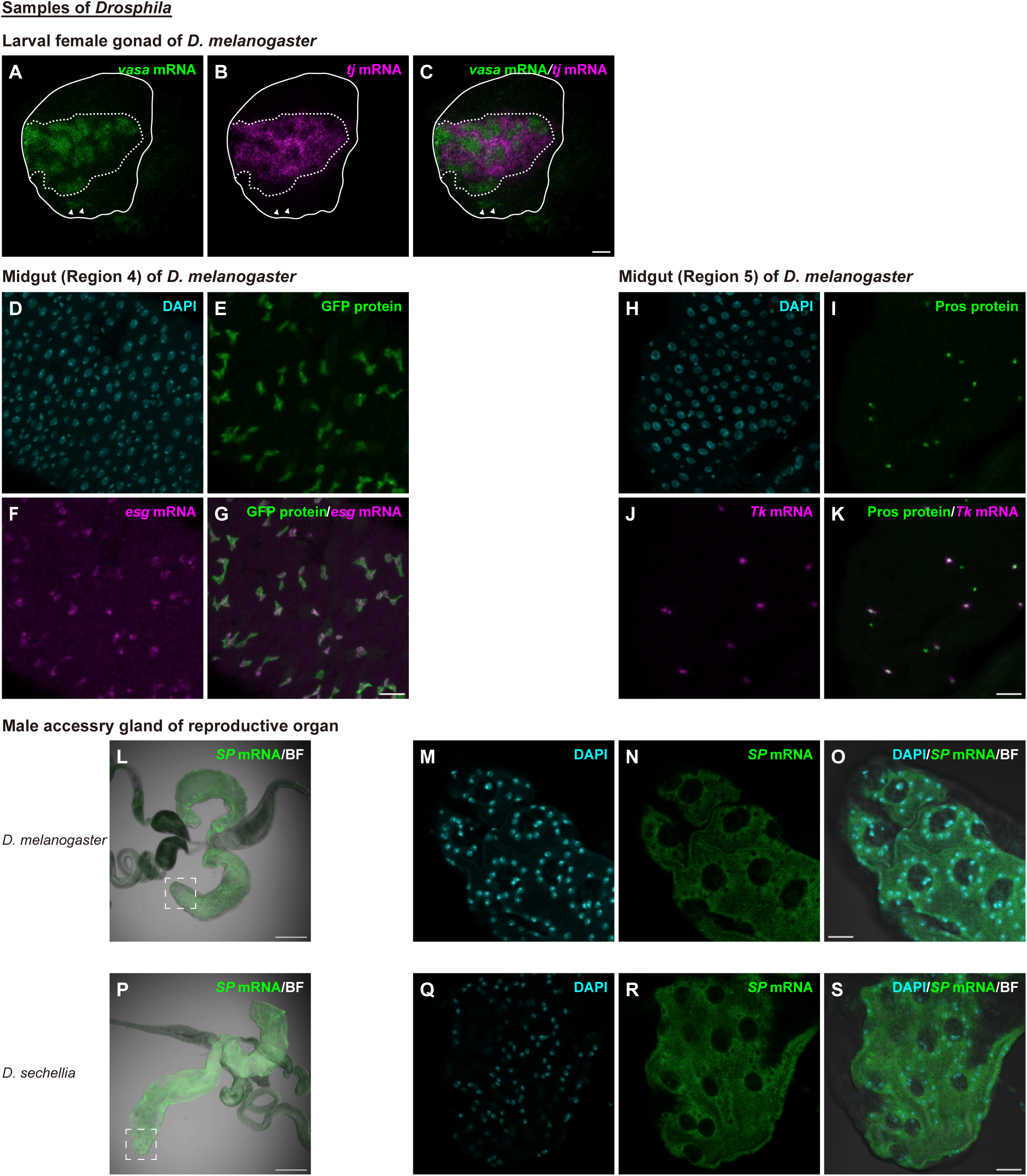
Applicability of EC-isHCR in various whole-mount *Drosophila* samples. (A–C) Confocal microscopy images of third-instar larval ovary from *D. melanogaster*. Green and magenta indicate *vasa* mRNA and *tj* mRNA, respectively. Merged image of *vasa* mRNA and *tj* mRNA signals is shown in (C). White lines and dotted lines outline the whole ovary and germline/ICs region, respectively. Arrowheads indicate germline cells located in the posterior region. Ovaries are oriented with anterior up, posterior down, medial left, and lateral right. Scale bar: 20 µm. (D–G) Confocal microscopy images of “Region 4” of the *D. melanogaster* midgut. Cyan, green, and magenta indicate DAPI, GFP driven by *esg*-GAL4, and *esg* mRNA signals, respectively. Merged image of GFP and *esg* mRNA signals is shown in (G). Scale bar: 20 µm. (H–K) Confocal microscopy images of “Region 5” of the *D. melanogaster* midgut. Cyan, green, and magenta indicate DAPI, Pros protein, and *Tk* mRNA signals, respectively. Merged image of Pros protein and *Tk* mRNA is shown in (G). Scale bar: 20 µm. (L–O) Bright-field (BF) and confocal microscopy images of male accessory glands from *D. melanogaster*. Cyan and green indicate DAPI and *SP* mRNA signals, respectively. Merged BF image with *SP* mRNA signal is shown in (L). A higher-magnification view of the region boxed in (L) is shown in (M–O). Merged image of DAPI and *SP* mRNA signals is shown in (O). Scale bars: 100 µm (L) and 20 µm (M–O). (P–S) BF and confocal microscopy images of male accessory glands from *D. sechellia*. Cyan and green indicate DAPI and *SP* mRNA signals, respectively. Merged BF image with *SP* mRNA signal is shown in (P). A higher-magnification view of the region boxed in (P) is shown in (Q–S). Merged image of DAPI and *SP* mRNA signals is shown in (S). Scale bars: 100 µm (P) and 20 µm (Q–S).

### 2.2. Applicability of EC-isHCR in whole-mount wasp and aphid samples

To examine whether EC-isHCR can be applied to whole-mount samples other than *Drosophila*, we focused on two non-model insects, *Asobara japonica* (*A. japonica*) and the pea aphid *Acyrthosiphon pisum* (*A*. *pisum*).

*A. japonica* is a parasitoid wasp that parasitizes a wide range of *Drosophila* species and degrades host imaginal discs by injecting venom [17,18]. Imaginal disc degradation factor-1 (IDDF-1), an abundant protein in the venom, induces imaginal disc degradation [18]. EC-isHCR successfully detected *IDDF-1* mRNA in the venom gland of *A. japonica* (Fig. 2A, B), consistent with the distribution of IDDF-1 protein (Fig. S1). These findings demonstrate that EC-isHCR can visualize mRNAs in wasps.

**Fig. 2.**
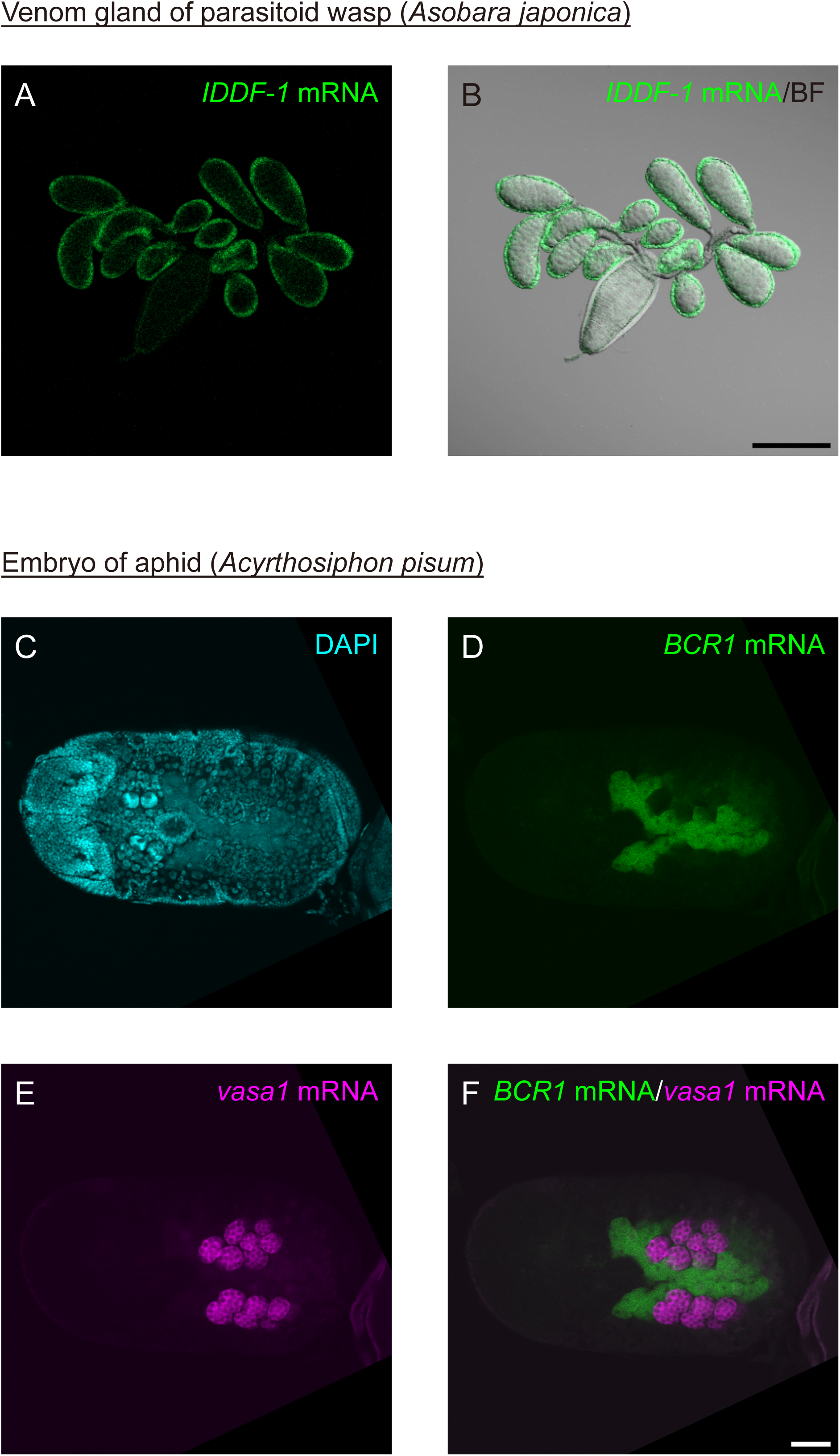
Applicability of EC-isHCR in whole-mount wasp and aphid samples. (A–B) BF and confocal microscopy images of venom glands from *A. japonica*. Green indicates *IDDF-1* mRNA signal. Merged BF image with *IDDF-1* mRNA signal is shown in (B). Scale bar: 200 µm. (C–F) Confocal microscopy images of a late *A. pisum* embryo after katatrepsis. Cyan, green, and magenta indicate DAPI, *BCR1*, and *vasa1* mRNA signals, respectively. Merged image of *BCR1* and *vasa1* mRNA signals is shown in (F). Scale bar: 50 µm.

*A. pisum* is a well-studied model of endosymbiosis. *A*. *pisum* possesses specialized host cells called bacteriocytes that harbor an obligate mutualistic bacterial endosymbiont, *Buchnera aphidicola* [19,20]. In these cells, *Bacteriocyte-specific Cysteine-Rich* (*BCR*) family genes, which encode antimicrobial peptides, are expressed throughout the life cycle from early embryogenesis to adult [21,22]. By contrast, expression of *vasa1* is germline-restricted and not observed in the bacteriocyte [23]. Consistent with previous reports [21–23], *BCR1* signals were specifically localized to bacteriocytes, while *vasa1* signals were detected in seven germ cell subgroups located bilaterally adjacent to the bacteriocytes (Fig. 2C–F). These findings indicate that EC-isHCR can also visualize mRNAs in aphids.

### 2.3. Applicability of EC-isHCR in paraffin trout sections

In the testis of rainbow trout (*Oncorhynchus mykiss: O. mykiss*), a salmonid species, germ cells are surrounded by Sertoli cells (a type of somatic cell). Germ cells and Sertoli cells express *vasa* [24] and *gonadal soma-derived growth factor* (*gsdf*) [25], respectively. Consistent with this, EC-isHCR detected *vasa* mRNAs enclosed by *gsdf* mRNA (Fig. 3A–C). These results indicated that EC-isHCR can be applied to sections of salmonid samples. Furthermore, because the modified version of third-generation isHCR (modified isHCR) has also been applied to salmonid samples [26], we compared signal intensities between EC-isHCR and modified isHCR. The signals obtained with EC-isHCR were equivalent to those observed in modified isHCR (Fig. 3). Because EC-isHCR and modified isHCR experiments require 1 day and 2 days, respectively, our results suggest that EC-isHCR can visualize mRNAs more rapidly than modified isHCR, without compromising signal intensity.

**Fig. 3.**
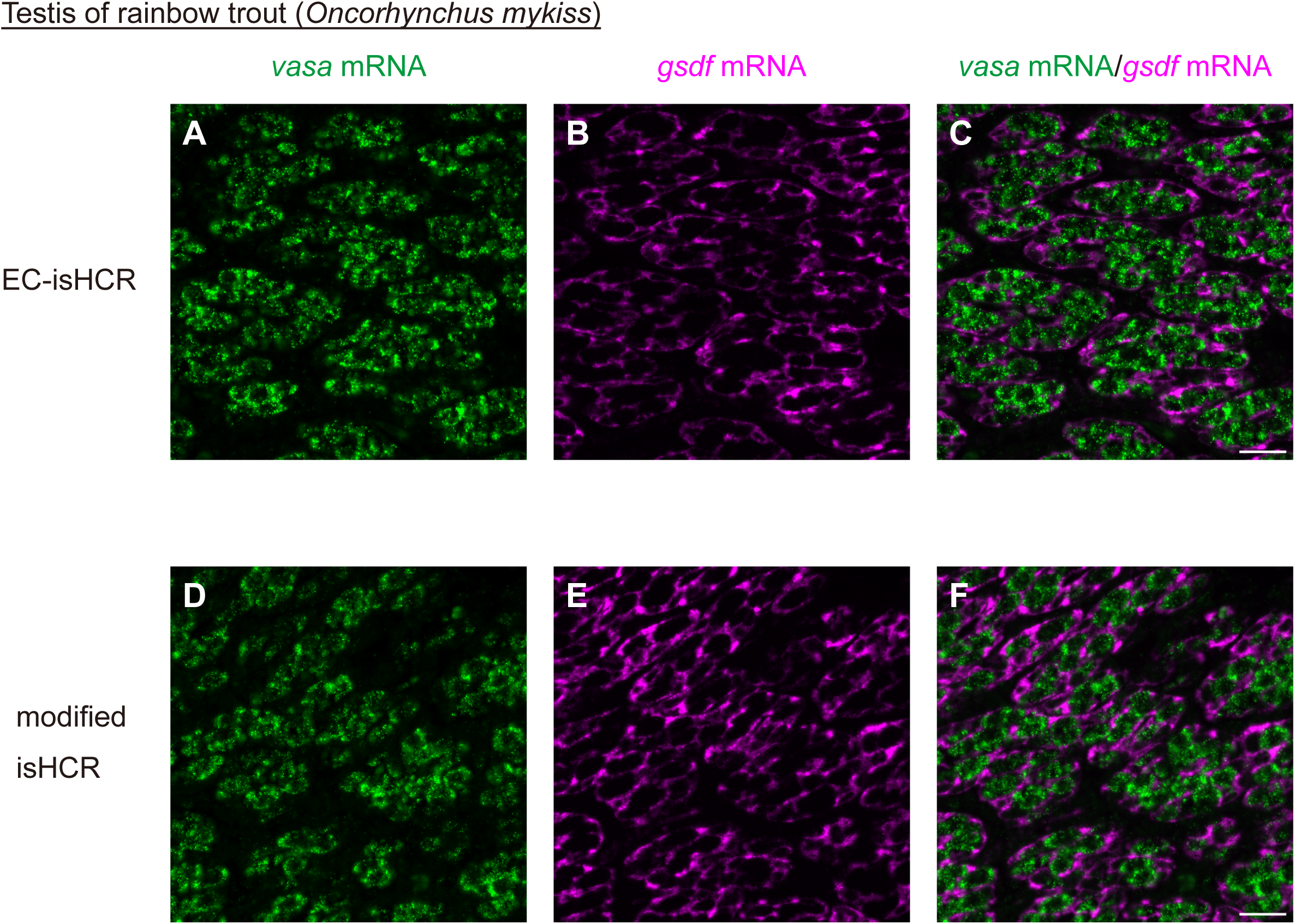
Applicability of EC-isHCR in paraffin trout sections. (A–F) Microscopy images of the testis sections from rainbow trout. Samples stained using EC-isHCR are shown in (A–C), and samples stained using modified isHCR are shown in (D–F). Green and magenta indicate *vasa* mRNA and *gsdf* mRNA signals, respectively. Merged images of *vasa* mRNA and *gsdf* mRNA signals are shown in (C) and (F). Scale bar: 20 µm.

### 2.4. Applicability of EC-isHCR in frozen mouse sections

We examined whether EC-isHCR can be applied to frozen mouse sections (*Mus musculus*: *M. musculus*). In the Allen Mouse Brain Atlas, the density of *glutamate decarboxylase 1* (*Gad1*) signals is high in the reticular formation (RF) and low in the striatum (STR) (https://connectivity.brain-map.org/transgenic/experiment/306566995) [27]. Consistent with this, EC-isHCR detected *Gad1* signals of high density in the RF and of low density in the STR (Fig. 4A, B, D). Similarly, as observed in the Allen Mouse Brain Atlas [27], EC-isHCR detected *preproenkephalin* (*Penk*) mRNA signals restricted to the STR (Fig. 4A, C, D). These results indicate that EC-isHCR is applicable to mouse brain sections.

**Fig. 4.**
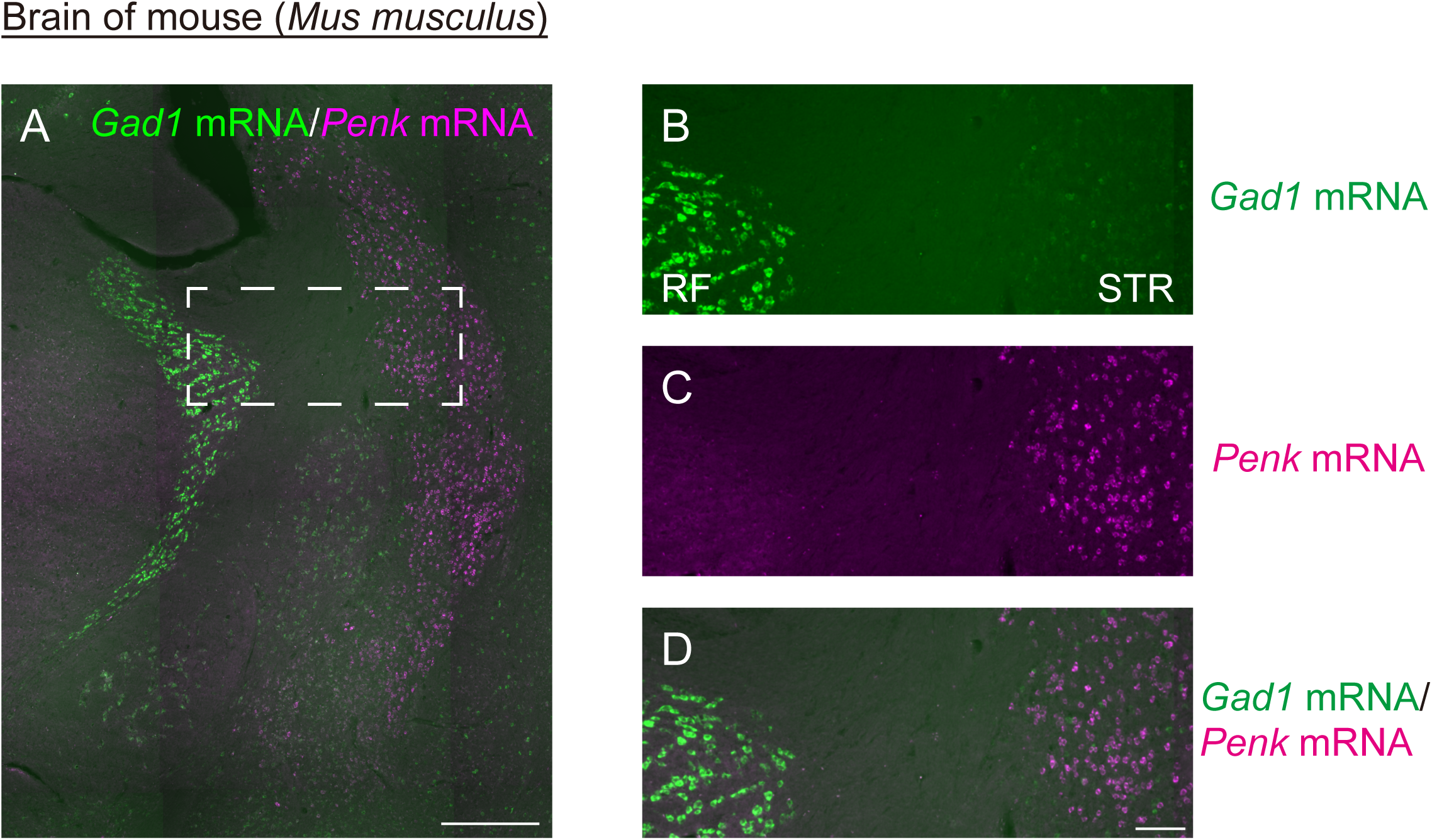
Applicability of EC-isHCR in frozen mouse sections. (A–D) Microscopy images of the brain section including reticular formation (RF) and striatum (STR). Green and magenta indicate *Gad1* mRNA and *Penk* mRNA signals, respectively. A higher-magnification view of the region boxed in (A) is shown in (B–D). Merged image of *Gad1* mRNA and *Penk* mRNA signals is shown in (D). Scale bars: 200 µm (A) and 50 µm (B–D).

### 2.5. EC-isHCR to visualize subcellular localization of mRNA

In addition to organ- and cell-level analyses, we examined whether EC-isHCR can detect mRNAs at the subcellular level. Germ granules are germline-specific RNP granules, which are membraneless organelles formed through the condensation of mRNAs and proteins [28,29]. In primordial germ cells of *Drosophila* embryos, germ granules contain *nanos* mRNA but not *Hsp83* mRNA [30,31]. Using EC-isHCR, we successfully detected *nanos* mRNA in germ granules marked with Vasa protein (Fig. 5A–C), whereas *Hsp83* mRNA was not localized in the granules (Fig. 5A, D, E). We also applied EC-isHCR to human cultured cells. *kinesin family member 1C* (*KIF1C*) mRNA is enriched in protrusions of HeLa cells [32,33], and EC-isHCR detected *KIF1C* mRNA in HeLa cell protrusions (Fig. 5F, G). Together, these results indicate that EC-isHCR visualizes subcellular localization of mRNA.

**Fig. 5.**
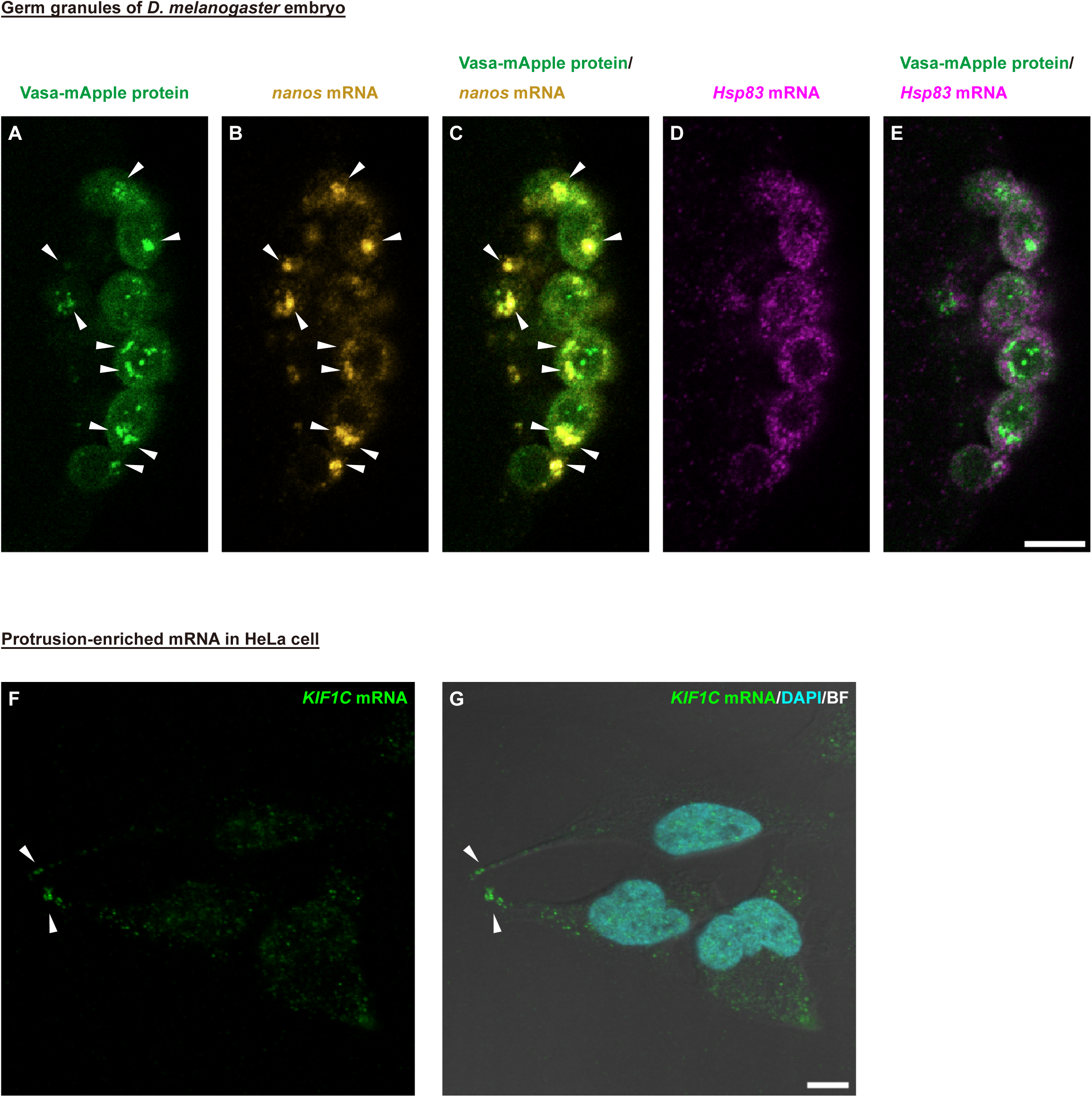
EC-isHCR to visualize subcellular localization of mRNA. (A–E) Confocal microscopy images of a blastodermal (nuclear-cycle-14) embryo from *D. melanogaster*. Green, yellow, and magenta indicate Vasa-mApple, *nanos* mRNA, and *Hsp83* mRNA signals, respectively. Merged image of Vasa-mApple with *nanos* mRNA and Vasa-mApple with *Hsp83* mRNA are shown in (C) and (E), respectively. Arrowheads indicate regions where Vasa-mApple and *nanos* mRNA signals are co-localized. Scale bar: 10 µm. (F, G) BF and confocal microscopy images of HeLa cells. Green and cyan indicate *KIF1C* mRNA and DAPI signals, respectively. Merged image of the *KIF1C* mRNA signal, DAPI staining, and BF image is shown in (G). Arrowheads indicate protrusions where *KIF1C* mRNA signals are enriched. Scale bar: 10 µm.

## 3. Discussion

In this study, we demonstrated that EC-isHCR detected mRNAs across a broad range of samples, such as whole-mount preparations of fruit flies, parasitoid wasps, and aphids; paraffin sections of trout; frozen mouse sections; and human cultured cells. Our results suggest that EC-isHCR could be applicable to a broad range of samples prepared using various methods, as described above.

EC-isHCR visualized mRNAs at the organ level (*Drosophila* MAG, wasp venom gland, and mouse brain), cellular level (*Drosophila* larval gonad and gut, aphid bacteriocytes, rainbow trout testis), and subcellular level (*Drosophila* germ granules and HeLa cell protrusions). Thus, EC-isHCR detected RNAs across multiple spatial scales, which is a key feature of conventional isHCR.

EC-isHCR obtained a signal intensity comparable to modified isHCR (Fig. 3). Furthermore, EC-isHCR successfully visualized various mRNAs that were also detected by either third-generation isHCR or modified isHCR in previous reports [4,5,13]. Therefore, these results suggest that EC-isHCR does not compromise signal intensity and retains high sensitivity for RNA detection.

In this study, we additionally developed an automated probe design tool for EC-isHCR (Supplemental Information 1, https://github.com/ShuntaYorimoto/hcrkit). Although we previously released a probe design tool [8], it had several limitations. For example, off-target evaluation with BLAST had to be performed manually, which limited the usability of the tool. To overcome this, we integrated a local BLAST module into the probe design workflow, enabling automatic off-target evaluation. Furthermore, we improved the tool to increase the number of probes. The previous version of our tool identified candidate regions for probe hybridization based on GC content, avoiding overlaps, and then off-target evaluation had to be performed with BLAST. In the previous workflow, overlapping candidate regions were eliminated before off-target evaluation, resulting in the exclusion of candidates that meet the off-target evaluation criteria. We modified the tool to first detect candidate regions while permitting overlaps, perform BLAST-based evaluation, and then select non-overlapping regions for probes. These modifications enhanced the probe design tool and broadened the applicability of EC-isHCR. By combining the tool with EC-isHCR, our study establishes a framework that reduces the barrier to using fast isHCR.

Our framework can make the usability of the isHCR comparable to, or even higher than other ISHs. For example, RNAscope is often used for tissue or cell-level analyses. RNAscope, which visualizes mRNA through probe hybridization followed by some rounds of amplification [6], like isHCR, is used to analyze RNA expression patterns in samples[34–36], validate single-cell RNA-seq analyses [4,37,38], or assess gene expression changes induced by genetic tools [39–42]. RNAscope can complete staining within a single day [6] while third-generation isHCR and modified isHCR require 3 days and 2 days for staining, respectively. Thus, conventional isHCR is less efficient in terms of staining time than RNAscope. Our fast EC-isHCR achieved staining times comparable to those of RNAscope, thereby addressing this shortcoming. Furthermore, our probe design tool is openly accessible, whereas the manufacturer of RNAscope does not disclose the precise probe sequences [34]. Thus, our framework makes isHCR a more accessible and appealing option. Subcellular analysis is commonly conducted by smFISH [43–46]. smFISH visualizes RNA with probes directly conjugated to fluorophores [47,48] or with probes hybridized to fluorophore-labeled oligonucleotides [7]. Because smFISH does not employ signal amplification, signal intensity might be modest, potentially making smFISH unsuitable for low-copy RNAs. HCR, the amplification reaction, does not rapidly reach a reaction plateau [49,50]. Consequently, extending the reaction time can improve the signal-to-noise ratio and the detection of low-copy RNAs. Taken together, our fast EC-isHCR is a practical option to visualize subcellular localization of RNAs.

Our framework can contribute to a wide range of research applications. EC-isHCR requires transcriptomic data and, ideally, genomic data to design target-specific probes. Such resources are becoming increasingly available: transcriptomic and whole-genome sequence data are rapidly accumulating for many non-model animals [51–53]. Consistent with this, EC-isHCR was successfully applied to animals with available transcriptomic and genomic data, such as *D. sechellia*, trout, wasp, and aphid, and this study reports the first application of isHCR to wasps and aphids. Therefore, our framework promotes the applicability of isHCR in basic studies of ecology and evolutionary biology, as well as in applied approaches in medicine and agricultural science.

In conclusion, EC-isHCR shortens staining time while retaining the key features of conventional isHCR, providing an attractive option for RNA detection compared with other ISH methods. Furthermore, by combining EC-isHCR with the automated probe design tool, we provide a framework that lowers the barrier to use. This platform is expected to serve as a powerful driver for research across diverse fields of life science.

## 4. Materials and methods

### 4.1. Split-initiator DNA probes and hairpin DNAs

The split-initiator DNA probes were designed as described previously [5,34,54]. Candidate probe binding sites were selected with a GC content of 40–60%. Sites with less than 50% sequence homology to non-target RNAs, as determined by BLAST, were retained. Their reverse complement sequences were conjugated with partial initiator sequences. Probe sequences for each gene are listed in Table S1. We developed a custom tool to automate the entire probe design workflow, which is available on GitHub (https://github.com/ShuntaYorimoto/hcrkit).

We selected hairpin DNAs, including the initiators (the toehold and stem domains of hairpin H2), following the manufacturer’s instructions and a previous study [26]. For model organisms and cultured cells, we used hairpin DNAs validated by the manufacturer. For non-model organisms, candidates were examined according to the manufacturer’s instructions. Briefly, we used BLAST to confirm ≤13 nucleotides of initiator sequence identity against reference RNA sequences; if this was not satisfied, we checked for ≤4 nucleotides within the toehold domain. When hairpin DNAs did not meet these thresholds, we tested their usability in the target sample, as in the previous study [26], by comparing signals between experimental and control groups (Fig. S2). The hairpin DNAs for each sample were listed in Table S2.

### 4.2. Animal husbandry and cell culture

For the experiments of *Drosophila* embryos and larval gonads, flies were maintained on standard *Drosophila* medium at 25°C. For the experiments of *Drosophila* guts and MAGs, flies were maintained on standard *Drosophila* medium at 18°C or 25°C in a 12-h:12-h light/dark cycle.

*A. japonica* were reared on standard agar-cornmeal medium at 25°C in a 12-h:12-h light/dark cycle.

*A. pisum* were maintained on vetch seedlings (*Vicia faba*) in growth chambers at 16°C in a 16-h light:8-h dark cycle.

All fish procedures were approved by the Institutional Animal Care and Use Committee, Tokyo University of Marine Science and Technology. Rainbow trout were maintained at 10°C at the Tokyo University of Marine Science and Technology, Oizumi Station (Yamanashi, Japan).

All mouse experiments were approved by the Institutional Animal Care and Use Committee of the University of Tsukuba, and all experiment procedures were conducted in accordance with the Guidelines for Animal Experiments of the University of Tsukuba. Mice were housed in standard cages and maintained under a controlled environment (23.5 ± 2.0°C, 50.0 ± 10.0% humidity, 12/12 h light/dark cycle, lights on at 9 A.M.). Food and water were available *ad libitum*.

HeLa cells were maintained in Dulbecco’s Modified Eagle Medium (FUJIFILM Wako) supplemented with 10% fetal bovine serum (Gibco, Cat#10270-106), 100 U/mL penicillin/streptomycin (Gibco, Cat#15140122) at 37°C with 5% CO2.

### 4.3. Genotype and sample condition

For experiments using *Drosophila* samples, *Oregon-R*, *w*, *y w*, *w*; *vasa-mApple* [55] (Kyoto *Drosophila* Stock Center; stock# 118617, RRID: DGGR_118617) and *w; esg-GAL4, UAS-GFP, tub-GAL80ts/CyO* [56] (RRID:BDSC_93857, gift from Fumiaki Obata) were used. For the experiments using *O. mykiss* samples, WT and p*vasa*-*GFP* transgenic rainbow trout [57,58] were used. Strains of *D. sechellia*, *A.japonica*, *A. pisum*, and *M. musculus* were *3C* (KYORIN-Fly: *Drosophila* species stock center; stock# k-s10), *Tokyo (TK)* [59]*, ApL*, and C57BL/6J, respectively. Detailed sample conditions are provided in Table S3.

### 4.4. Histology and sample fixation

Compositions of phosphate-buffered saline (PBS) and saline-sodium citrate buffer (SSC) used for each sample type are listed in Table S4.

*Drosophila* embryos were collected on grape juice agar plates and dechorionated in sodium hypochlorite solution. Following dechorionation, embryos were fixed in a 1:1 mixture of heptane and fixative (4% formaldehyde in PBS) for 30 min. Fixed embryos were shaken vigorously in 1:1 MeOH:heptane to remove vitelline membranes. The embryos were then rinsed three times with MeOH and stored in MeOH at −20°C. Before EC-isHCR, samples were dehydrated with 3:1, 1:1, and 1:3 MeOH:1× SSCT (SSC with 0.1% Tween-20).

*Drosophila* larval gonads were dissected in Ephrussi–Beadle Ringer solution (130 mM NaCl, 5mM KCl, 2mM CaCl_2_, 10mM HEPES pH 6.9) [60] and fixed in 6% formaldehyde in buffer B (16.7mM KH_2_PO_4_/K_2_HPO_4_, 75mM KCl, 25 mM NaCl, 3.3 mM MgCl_2_, pH 6.8) for 10 min at room temperature (RT). The fixed ovaries were immediately washed twice with 0.2% PBTw (0.2% Tween-20 in 1× PBS) for 30 sec, once for 3 min and once for 10 min once. Samples were sequentially rinsed with 3:1, 1:1, and 1:3 0.2% PBTw:1× SSCT.

*Drosophila* adult samples were dissected in cold 1× PBS and fixed in 4% formaldehyde (Nacalai Tesque, Cat# 02890-45) in PBS for 20 to 30 min (MAGs) or 30 to 40 min (guts) at RT. Fixed samples were washed three times in 0.1% PBTr (0.1% Triton X-100 in 1× PBS). Samples were sequentially dehydrated in 1:3, 1:1, and 3:1 mixtures of either PBS:MeOH (MAGs) or 0.1% PBTr:MeOH (guts), and placed in 100% MeOH overnight at −20°C (MAGs) or for 20 min at RT (guts). Samples were sequentially rinsed with 3:1, 1:1, and 1:3 MeOH:1× SSCT.

*A. japonica* venom glands were dissected in PBS (Nacalai Tesque, Cat# 27575-31) and fixed with 3.7% formaldehyde in 0.3% PBTr (10-fold dilution of a 37% formaldehyde solution, Nacalai Tesque, Cat# 16222-65) for 30 min at 25°C. Fixed samples were rinsed three times with 0.3% PBTr. Samples were sequentially rinsed with 3:1, 1:1, and 1:3 0.3% PBTr:1× SSCT. Notably, these sequential rinses were performed only for EC-isHCR, not for immunohistochemistry.

Viviparous ovaries of *A*. *pisum* were dissected from 4th instar nymphs or young adults in 4% PFA in PBS (FUJIFILM Wako Pure Chemical Corporation, Cat# 163-20145) and fixed with fresh 4% PFA in PBS for 30 min at RT. The fixed samples were dehydrated with a 1:1 mixture of MeOH and water (v:v) for 30 min and stored in 100% MeOH at −20°C until use. The samples were sequentially rinsed with 3:1, 1:1, and 1:3 MeOH: 1× SSCT.

*O. mykiss* testes were fixed at 4°C for 16 h in 4% formaldehyde (FUJIFILM Wako Pure Chemical Corporation, Cat# 162-16065) in PBS. The fixed samples were dehydrated using a standard EtOH-xylene series and embedded in paraffin wax. The paraffin block was sliced into 4-µm serial sections. The paraffin sections were dewaxed and rehydrated by passing them through a xylene-EtOH series. After rehydration, the sections were incubated with MeOH at RT for 10 min, and rehydrated in 3:1, 1:1, 1:3 MeOH: 1× SSCT solutions for 2 min each.

*M. musculus* were deeply anesthetized and transcardially perfused with saline followed by 10% formalin neutral buffer solution (FUJIFILM Wako Pure Chemical Corporation, Cat# 068-01663). The brains were removed and postfixed in 10% formalin neutral buffer solution at 4°C overnight, and then incubated in 30% sucrose (w/v) in PBS at 4°C for 2 nights. The brains were embedded and frozen in OCT compound (Sakura Finetek, Cat#4583), and sectioned at 30 μm thick using a sliding microtome (Yamato Kohki, REM-710). Sections were washed with PBS for 5 min. Then sections were incubated with MeOH for 10 min. Samples were sequentially rinsed with 3:1, 1:1, and 1:3 MeOH:1× SSCT.

HeLa cells grown on glass coverslips were rinsed three times with PBS and fixed with 4% formaldehyde in PBS for 15 minutes. Fixed cells were rinsed three times with PBS and dehydrated in 70% EtOH. Samples were stored at −20°C overnight (up to 1 week).

### 4.5. EC-isHCR

EC-isHCR was performed as described previously [8] with minor modifications for sample conditions. For HeLa cells, the rinse and wash steps with 1× SSCT in the following procedures were replaced with 1× SSC. The samples were washed three times with 1× SSCT, and then pre-hybridized in pre-warmed hybridization buffer [15% ethylene carbonate (Sigma-Aldrich, Cat# E26258), 5× SSC, 0.1% Tween-20, 50 μg/ml heparin (Sigma-Aldrich, Cat# H3393-25KU), 1× Denhardt’s solution (Fujifilm-Wako, Cat# 043-21871)] for 30 min at 45°C. Probe sets for each mRNA were diluted in hybridization buffer at 20 nM and the samples were incubated with this probe solution for 2 h at 45°C. After hybridization, the samples were washed twice with hybridization buffer for 10 min at 45°C. The samples were rinsed with 1:1 hybridization buffer: once with 1× SSCT and then twice with 1× SSCT, and then washed with 1× SSCT for 10 min at RT. Before the amplification, hairpin DNA pairs (H1, H2) were snap-cooled; each hairpin DNA was placed in a separate tube and warmed at 95°C for 2 min, and then cooled to 65°C for 15 min and to 25°C for 40 min. The hairpin DNA pairs were diluted 1:50 in amplification buffer (Nepagene, Cat# IPL-AB) to make the amplification solution. The samples were incubated with the amplification solutions for 2 h at 25°C. After the amplification, the samples were washed three times with 1× SSCT at RT and mounted. To stain MAG nuclei, 4’,6-diamidino-2-phenylindole (DAPI; Thermo Fisher Scientific, Cat# 62247; 1:10,000) was added during the final wash. The samples were then rinsed once with 1× SSCT and mounted. The mounting and observation conditions are shown in Table S5.

### 4.6. modified isHCR

Modified isHCR was performed according to the previous report [26] and manufacturer’s instructions with minor modifications. After rehydration, the sections were treated with MeOH at RT for 10 min, washed twice with 0.1% PBTw for 5 min each, and then equilibrated with hybridization buffer (Nepagene, Cat# IPL-HB) in a moist chamber at 37°C for 30 min. The probe mixture containing 20 nM of each probe in hybridization buffer was heat-denatured at 95°C for 3 min. The mixture was applied to the sections and incubated at 37°C overnight in a moist chamber for hybridization. After hybridization, the slides were washed three times with 0.5× SSCT at 37°C for 10 min. Hairpin DNA pairs were snap-cooled as in EC-isHCR and were diluted 1:50 in amplification buffer (Nepagene, Cat# IPL-AB) to make the amplification solution. The samples were incubated with the amplification solutions for 2 h at 25°C. The slides were washed three times with 0.5× SSCT at 37°C for 10 min. The mounting and observation conditions were the same as for EC-isHCR.

### 4.7. Immunohistochemistry

After EC-isHCR amplification, *Drosophila* guts were rinsed with 1× SSCT, then with 1:1 1× SSCT: 0.1% PBTr, and finally three times with 0.1% PBTr, and incubated for 20 min in 0.1% PBTr. Samples were then incubated in blocking solution [0.1% PBTr with 2% BSA (Sigma-Aldrich, Cat# A9647-1006)] for 1 h at RT, and then incubated overnight with blocking solution containing mouse anti-Prospero (1:100; Developmental Studies Hybridoma Bank, Clone ID: MR1A, RRID: AB_528440) at 4°C. The next day, the solution of the primary antibodies were removed, rinse three rinses with 0.1% PBTr, and then incubated for 20 min in 0.1% PBTr. Samples were then incubated for 2 h at RT, with Alexa Fluor 555-conjugated secondary antibodies (1:200; Thermo Fisher Scientific, Cat# A32727, RRID: AB_2633276) in a blocking solution. Samples were incubated three times in 0.1% PBTr, for 20 min each. During the last wash, DAPI (1:10,000) was added to visualize the nucleus. Samples were finally mounted and observed as shown in Table S4.

Fixed venom glands of *A. japonica* were blocked with blocking solution (2% BSA in 0.3% PBTr) for 90 min at 25°C followed by a treatment with the primary antibody (anti-IDDF-1 rabbit antibody [18], RRID: AB_3717376,1:100 in blocking solution) at 4°C overnight. Samples were washed three times with 0.3% PBTr and incubated with Alexa Fluor 488-conjugated secondary antibody (anti-rabbit IgG, Thermo Fisher Scientific, Cat# A32731, RRID: AB_2633280) and phalloidin (Thermo Fisher Scientific, Cat# A34055) at 25°C for 2.5 h. After washing three times with 0.3% PBTr, the samples were mounted and observed as in Table S5.

### 4.8. Images

Minor modification of rotation and linear brightness/contrast adjustments were performed with Fiji (RRID: SCR_002285) [61].

## Acknowledgements

This work was supported by following grants; Grants-in-Aid for Scientific Research from the Japan Society for the Promotion of Science (JSPS) [KAKENHI Grant Numbers: 24H02030 (SK), JP23KJ0283 (HO), 23K05778 (MA), 23K26997 (MH) and 24H02060 (YWI)], Japan Agency for Medical Research and Development (AMED) [Grant number: JP24zf0127011 (YH)], and Japan Science and Technology Agency (JST) [Support for Pioneering Research Initiated by the Next Generation (SPRING) Grant Numbers: JPMJSP2124 (YK)].

## Disclosure statement

No potential conflict of interest was reported by the author(s).

## Author contributions

Conceptualization, Y.K. and K.M.; Data curation, Y.K., K.M., S.Y., T.H., H.O., Q.Q., R.H., T.K., Y.S., M.A., S.O., M.H., and S.S.; Formal analysis, Y.K., K.M., S.Y., T.H., H.O., Q.Q., R.H., T.K., Y.S., M.A., S.O., M.H., and S.S.; Funding acquisition, Y.K., H.O., M.H., Y.H., and S.K.; Investigation, Y.K., K.M., S.Y., T.H., H.O., Q.Q., R.H., T.K., Y.S., M.A., S.O., S.A., and M.H.; Methodology, Y.K., K.M., S.Y., T.H., H.O., Q.Q., R.H., T.K., Y.S., M.A., S.A., M.H., Y.H., and R.N.; Project administration, Y.K., K.M., S.Y., T.H., H.O., Q.Q., R.H., T.K., Y.S., M.A., M.H., Y.H., S.S., R.N., and S.K.; Resource, Y.K., Y.W.I., M.H., Y.H., S.S., R.N. and S.K.; Software, S.Y.; Supervision, Y.K., K.M., S.Y., R.N., and S.K.; Validation, Y.K., K.M., S.Y., M.H., Y.H., S.S., R.N., and S.K.; Writing – original draft, Y.K. and K.M.; Writing – review and editing, Y.K., K.M., S.Y., T.H., H.O., Q.Q., R.H., T.K., Y.S., M.A., S.O., S.A., Y.W.I., M.H., Y.H., S.S., R.N., and S.K.;

## Notes

https://github.com/ShuntaYorimoto/hcrkit

### Competing Interest Statement

The authors have declared no competing interest.

