## Supplemental Figures for "EC-isHCR: a rapid method for *in situ* hybridization chain reaction in diverse animal samples"

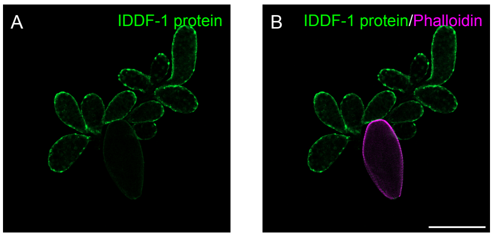


**Figure S1. Immunostaining for IDDF-1 protein in *A. japonica* venom gland.**

(A, B) Confocal microscopy images of an *A. japonica* venom gland. Green and magenta indicate IDDF-1 protein and phalloidin signals, respectively. Merged image of IDDF-1 and phalloidin signals is shown in (B). Scale bar: 200 µm.

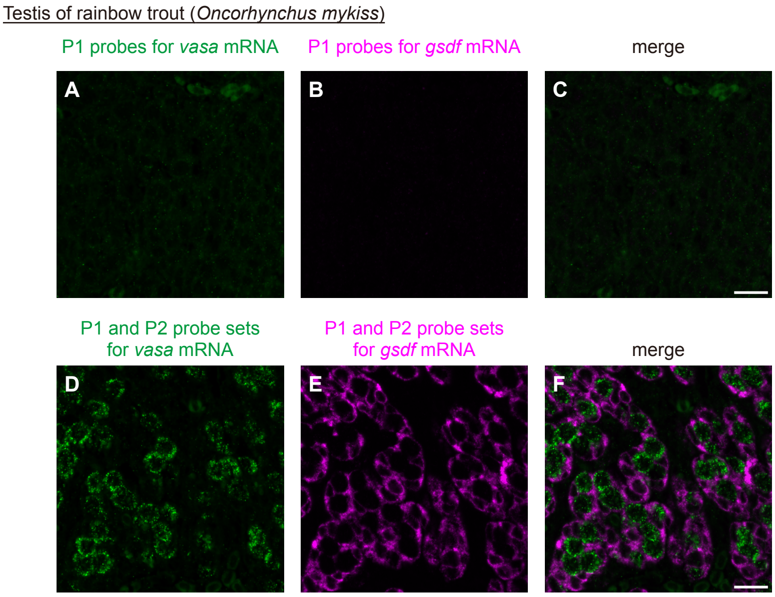


**Figure S2. Hairpin DNAs were available in sections of trout testis.**

(A–F) Microscopy images of testis sections from rainbow trout using the modified isHCR. Samples stained with S41 and A161 hairpin DNAs are shown in green (A, D) and magenta (B, E), respectively. For hybridization, *vasa* mRNA probes were used in (A, D), and *gsdf* mRNA probes were used in (B, E). Only P1 probes were used in (A, B) as controls, and both P1 and P2 probe sets were used in (D, E) as experiments. Although S41 and A161 hairpin DNAs with P1 probes alone provided weak or no signals (A, B), these hairpin DNAs produced strong signals with both the P1 and P2 probe sets (D, E), indicating that these hairpin DNAs are applicable for rainbow trout testis. Merged images of control and experimental groups are shown in (C) and (F). Scale bars: 20 µm.
