## Supplemental Information for "EC-isHCR: a rapid method for *in situ* hybridization chain reaction in diverse animal samples"

ShuntaYorimoto / hcrkit
Code Issues Pull requests Actions Projects Security
main hcrkit / OVERVIEW.md Go to file t
ShuntaYorimoto Update OVERVIEW.md 3b27979 · 2 days ago
171 lines (112 loc) · 7.75 KB
Preview Code Blame Raw

### Supplemental Information


- What is isHCR (*in situ* Hybridization Chain Reaction)?
  - Guidline for design of probes
- Overview of `hcrkit`
  - How does `hcrkit` remove probe regions based on off-target BLAST matches?
    - 1. if there are no isoforms other than the target transcript (default)
    - 2. if there are isoforms in addition to the target transcript
- Workflow & algorithm
  - Detect candidate probe regions based on GC content
  - Remove probe regions with off-target BLAST matches
    - BLAST Search
    - Filter by Specificity
  - Perform several processes to generate final probes
    - Select Non-overlapping Probe Regions
    - Generate Probe Sets
  - Write Outputs


#### What is isHCR (*in situ* Hybridization Chain Reaction)?

*in situ* hybridization chain reaction (isHCR), a type of *in situ* hybridization (ISH), enables RNA detection across multiple spatial scales, from organs to subcellular structures. Unlike enzyme-based methods, isHCR visualize RNAs though formation of polymers composed of fluorescently-labeled oligonucleotides.
The probe contains an initiator sequence (Figure SI1A). The initiator hybridizes with hairpin DNAs, triggering a hybridization chain reaction that produces fluorescently labeled polymers (Figure SI1B). To suppress background signals, the probe is split (P1 and P2), and the probe set can efficiently trigger amplification.

##### Guidline for design of probes

- GC content: 40–60% (45–55% recomended)
- For each probe set, sequence identity with off-target RNAs: ≤ 50% (verify using BLAST)
- Design region in target RNA: CDS + UTR (CDS recomended)

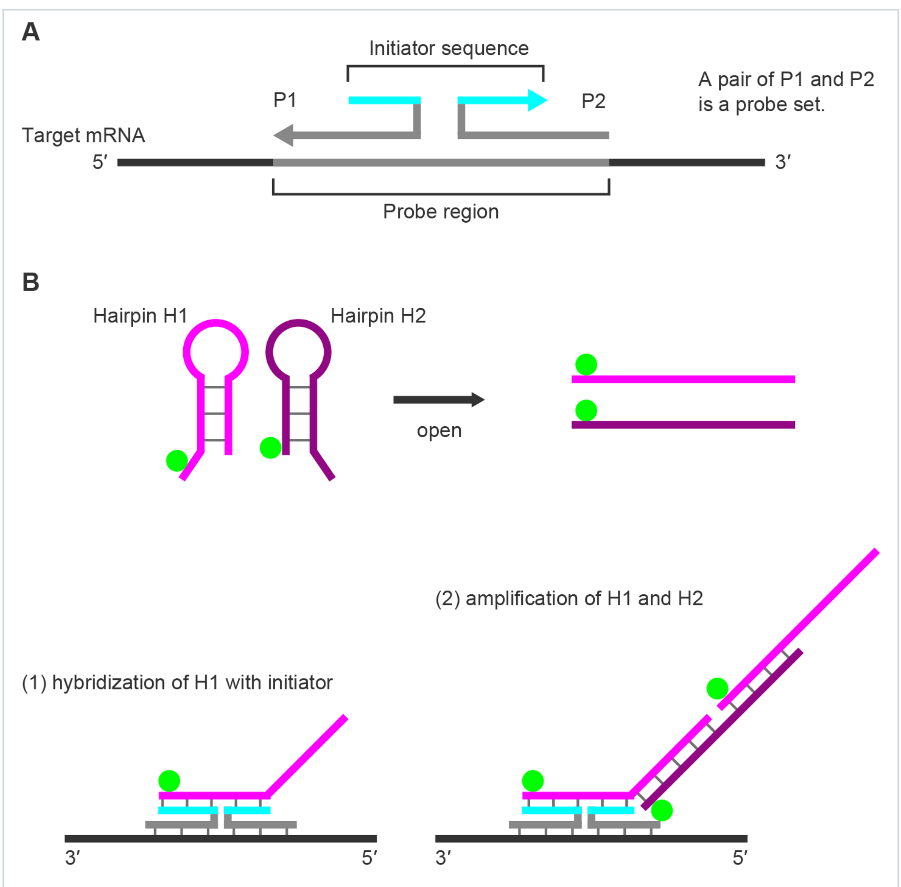


Figure SI1

#### Overview of `hcrkit`

`hcrkit` designs probes following the workflow below:

1. Detect candidate probe regions based on GC content
2. Remove probe regions based on off-target BLAST matches
3. Perform several processes to generate final probes

##### How does `hcrkit` remove probe regions based on off-target BLAST matches?

`hcrkit` excludes candidate probe regions that show BLAST hits to off-target sequences. Because BLAST also returns hits to the intended on-target sequences, these on-target hits must be ignored when evaluating potential off-target matches. To enable this filtering, `hcrkit` first obtains the IDs of on-target sequences using the three different methods described below (Figure SI4).

###### 1. If there are no isoforms other than the target transcript (default)

The on-target ID (Figure SI2, highlighted in green) is extracted from the FASTA header (the string before the first whitespace).

Example:

```
>XM_003242475.4 PREDICTED: Acyrthosiphon pisum nanos-like protein (LOC100216490), transcript variant X3, mRNA
AAAAAAATGTATGTTTTGAATTTGAAGTTTAAATTTAAGTGGGTGTTTTCAAGAGGACCCTTATTGCCAAGATATCACAA
TGAGTGGCCGTAGCAGATTCGCGTTCGGCAGCCAGACCCAGGCGCCCCAGTACCAGAACCGATGCGCTCTGTGCTTGACC
...
```

Figure SI2

###### 2. If there are isoforms in addition to the target transcript

###### 2-A. Automatically find out on-target IDs (with options `--gff3` + `--gene_name`)

The on-target IDs are identified from the GFF file. `hcrkit` processes the file as follows (Figure SI3):

- Select rows with Feature = "mRNA" (highlighted in cyan)
- Match gene= value in Attributes to `--gene_name` (highlighted in magenta)
- Extract transcript IDs as on-target isoforms (highlighted in green)

Example:

```
NC_042494.1 Gnomon      gene  112769404    112776872    .     -     .     ID=gene-
LOC100216490;Dbxref=GeneID:100216490;Name=LOC100216490;gbkey=Gene;gene=LOC100216490;gene_biotype=protein_coding
NC_042494.1 Gnomon      mRNA  112769404    112776872    .     -     .     ID=rna-XM_016802089.2;Parent=gene-
LOC100216490;Dbxref=GeneID:100216490,GenBank:XM_016802089.2;Name=XM_016802089.2;gbkey=mRNA;gene=LOC100216490;mo
del_evidence=Supporting evidence includes similarity to: 1 mRNA%2C 1 EST%2C 2 Proteins%2C and 100%25 coverage
of the annotated genomic feature by RNAseq alignments%2C including 20 samples with support for all annotated
introns;product=nanos-like protein%2C transcript variant X2;transcript_id=XM_016802089.2
NC_042494.1 Gnomon      exon  112776643    112776872    .     -     .     ID=exon-XM_016802089.2-1;Parent=rna-
XM_016802089.2;Dbxref=GeneID:100216490,GenBank:XM_016802089.2;gbkey=mRNA;gene=LOC100216490;product=nanos-like
protein%2C transcript variant X2;transcript_id=XM_016802089.2
NC_042494.1 Gnomon      exon  112771497    112772028    .     -     .     ID=exon-XM_016802089.2-2;Parent=rna-
XM_016802089.2;Dbxref=GeneID:100216490,GenBank:XM_016802089.2;gbkey=mRNA;gene=LOC100216490;product=nanos-like
protein%2C transcript variant X2;transcript_id=XM_016802089.2
NC_042494.1 Gnomon      exon  112769404    112769700    .     -     .     ID=exon-XM_016802089.2-3;Parent=rna-
XM_016802089.2;Dbxref=GeneID:100216490,GenBank:XM_016802089.2;gbkey=mRNA;gene=LOC100216490;product=nanos-like
protein%2C transcript variant X2;transcript_id=XM_016802089.2
NC_042494.1 Gnomon      CDS   112771497    112772020    .     -     0     ID=cds-XP_016657578.1;Parent=rna-
XM_016802089.2;Dbxref=GeneID:100216490,GenBank:XP_016657578.1;Name=XP_016657578.1;gbkey=CDS;gene=LOC100216490;p
roduct=nanos homolog 1;protein_id=XP_016657578.1
NC_042494.1 Gnomon      CDS   112769577    112769700    .     -     1     ID=cds-XP_016657578.1;Parent=rna-
XM_016802089.2;Dbxref=GeneID:100216490,GenBank:XP_016657578.1;Name=XP_016657578.1;gbkey=CDS;gene=LOC100216490;p
roduct=nanos homolog 1;protein_id=XP_016657578.1
NC_042494.1 Gnomon      mRNA  112769404    112776872    .     -     .     ID=rna-XM_016802088.2;Parent=gene-
LOC100216490;Dbxref=GeneID:100216490,GenBank:XM_016802088.2;Name=XM_016802088.2;gbkey=mRNA;gene=LOC100216490;mo
del_evidence=Supporting evidence includes similarity to: 2 mRNAs%2C 3 ESTs%2C 2 Proteins%2C and 100%25 coverage
of the annotated genomic feature by RNAseq alignments%2C including 38 samples with support for all annotated
introns;product=nanos-like protein%2C transcript variant X1;transcript_id=XM_016802088.2
NC_042494.1 Gnomon      exon  112776643    112776872    .     -     .     ID=exon-XM_016802088.2-1;Parent=rna-
XM_016802088.2;Dbxref=GeneID:100216490,GenBank:XM_016802088.2;gbkey=mRNA;gene=LOC100216490;product=nanos-like
protein%2C transcript variant X1;transcript_id=XM_016802088.2
NC_042494.1 Gnomon      exon  112771497    112772031    .     -     .     ID=exon-XM_016802088.2-2;Parent=rna-
XM_016802088.2;Dbxref=GeneID:100216490,GenBank:XM_016802088.2;gbkey=mRNA;gene=LOC100216490;product=nanos-like
protein%2C transcript variant X1;transcript_id=XM_016802088.2
NC_042494.1 Gnomon      exon  112769404    112769700    .     -     .     ID=exon-XM_016802088.2-3;Parent=rna-
XM_016802088.2;Dbxref=GeneID:100216490,GenBank:XM_016802088.2;gbkey=mRNA;gene=LOC100216490;product=nanos-like
protein%2C transcript variant X1;transcript_id=XM_016802088.2
NC_042494.1 Gnomon      CDS   112771497    112772020    .     -     0     ID=cds-XP_016657577.1;Parent=rna-
XM_016802088.2;Dbxref=GeneID:100216490,GenBank:XP_016657577.1;Name=XP_016657577.1;gbkey=CDS;gene=LOC100216490;p
roduct=nanos homolog 1;protein_id=XP_016657577.1
NC_042494.1 Gnomon      CDS   112769577    112769700    .     -     1     ID=cds-XP_016657577.1;Parent=rna-
XM_016802088.2;Dbxref=GeneID:100216490,GenBank:XP_016657577.1;Name=XP_016657577.1;gbkey=CDS;gene=LOC100216490;p
roduct=nanos homolog 1;protein_id=XP_016657577.1
NC_042494.1 Gnomon      mRNA  112769404    112776254    .     -     .     ID=rna-XM_003242475.4;Parent=gene-
LOC100216490;Dbxref=GeneID:100216490,GenBank:XM_003242475.4;Name=XM_003242475.4;gbkey=mRNA;gene=LOC100216490;mo
del_evidence=Supporting evidence includes similarity to: 1 mRNA%2C 1 EST%2C 2 Proteins%2C and 100%25 coverage
of the annotated genomic feature by RNAseq alignments%2C including 12 samples with support for all annotated
introns;product=nanos-like protein%2C transcript variant X3;transcript_id=XM_003242475.4
NC_042494.1 Gnomon      exon  112776187    112776254    .     -     .     ID=exon-XM_003242475.4-1;Parent=rna-
XM_003242475.4;Dbxref=GeneID:100216490,GenBank:XM_003242475.4;gbkey=mRNA;gene=LOC100216490;product=nanos-like
protein%2C transcript variant X3;transcript_id=XM_003242475.4
NC_042494.1 Gnomon      exon  112771497    112772031    .     -     .     ID=exon-XM_003242475.4-2;Parent=rna-
XM_003242475.4;Dbxref=GeneID:100216490,GenBank:XM_003242475.4;gbkey=mRNA;gene=LOC100216490;product=nanos-like
protein%2C transcript variant X3;transcript_id=XM_003242475.4
NC_042494.1 Gnomon      exon  112769404    112769700    .     -     .     ID=exon-XM_003242475.4-3;Parent=rna-
XM_003242475.4;Dbxref=GeneID:100216490,GenBank:XM_003242475.4;gbkey=mRNA;gene=LOC100216490;product=nanos-like
protein%2C transcript variant X3;transcript_id=XM_003242475.4
NC_042494.1 Gnomon      CDS   112771497    112772020    .     -     0     ID=cds-XP_003242523.1;Parent=rna-
XM_003242475.4;Dbxref=GeneID:100216490,GenBank:XP_003242523.1;Name=XP_003242523.1;gbkey=CDS;gene=LOC100216490;p
roduct=nanos homolog 1;protein_id=XP_003242523.1
NC_042494.1 Gnomon      CDS   112769577    112769700    .     -     1     ID=cds-XP_003242523.1;Parent=rna-
XM_003242475.4;Dbxref=GeneID:100216490,GenBank:XP_003242523.1;Name=XP_003242523.1;gbkey=CDS;gene=LOC100216490;p
roduct=nanos homolog 1;protein_id=XP_003242523.1
```

Figure SI3

> **Note**
>
> When users specify `--gff3` + `--gene_name`, the list of on-target IDs is output as `{prefix}_target_ids.txt`

###### 2-B. Manually provide on-target IDs (with options `--target_ids`)

The on-target IDs are specified manually. It is useful when:

- Users know all of the on-target IDs beforehand.
- Users wish to edit the list of on-target IDs obtained via 2-A.

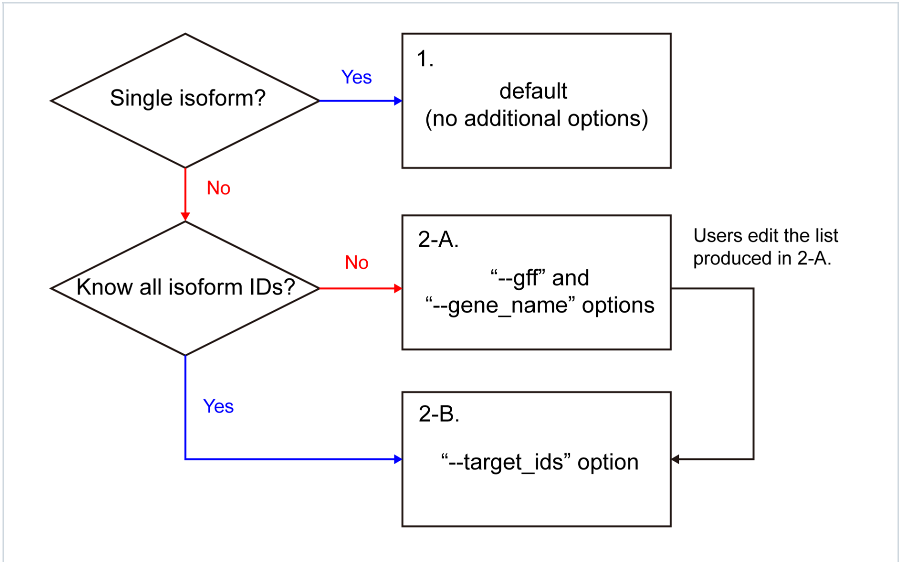


Figure SI4

#### Workflow & algorithm

`hcrkit` automates the probe design process through the following workflow:

##### Detect candidate probe regions based on GC content

###### Generate 52-nt sliding windows

The probe region for a split probe set is 52 nt. `hcrkit` divides the target transcript into overlapping 52-nt fragments using a sliding window (Figure SI5, moving 1 nt at a time from 5′ to 3′ end).

###### GC content filtering

Each 52-nt probe region is split into:

- P1 binding site: first 25 nt (positions 1-25)
- Spacer: 2 nt (positions 26-27)
- P2 binding site: last 25 nt (positions 28-52)

`hcrkit` calculates the GC content for both P1 and P2 binding sites. Only regions where **both** binding sites meet the GC criteria (default: 45–55%) are retained (Figure SI5).

###### Remove duplicates

If multiple probe regions have identical sequences, `hcrkit` keeps only one and removes the duplicates (Figure SI5).

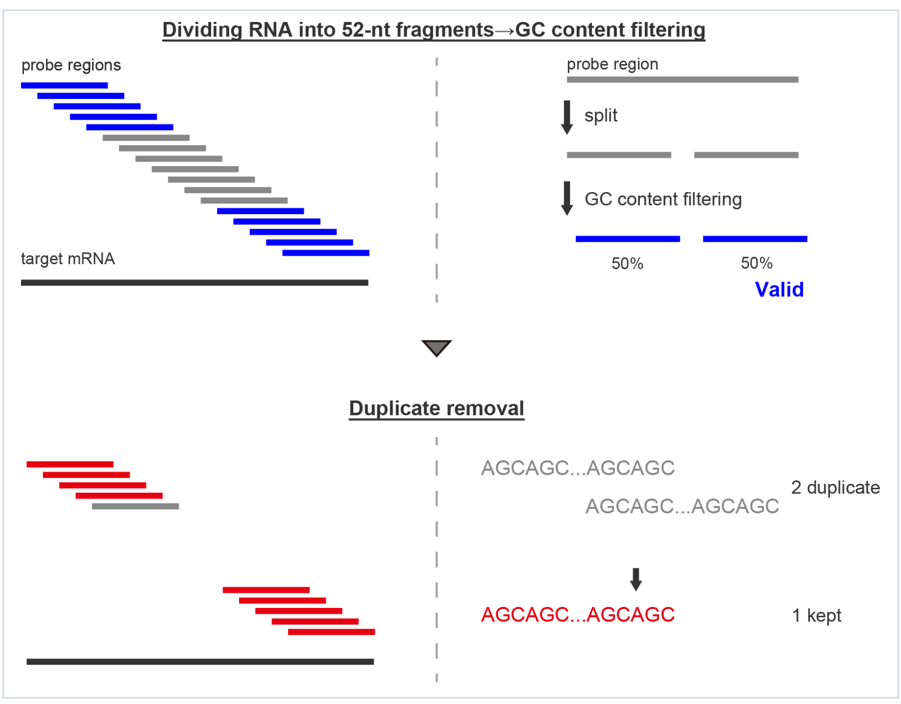


Figure SI5

##### Remove probe regions with off-target BLAST matches

###### BLAST Search

`hcrkit` performs BLASTN search against the reference transcriptome database to identify potential off-target binding sites for each probe region candidate (Figure SI6).

###### Filter by Specificity

For each BLAST hit, `hcrkit`:

1. Checks if the hit is to an on-target transcript
2. For off-target hits, calculates coverage: `(alignment length) / 52 * 100`
3. Tracks the maximum off-target coverage for each probe region

`hcrkit` removes probe regions where the maximum off-target coverage is ≥50% (Figure SI6). This ensures that each probe has sufficient specificity to the target transcript.

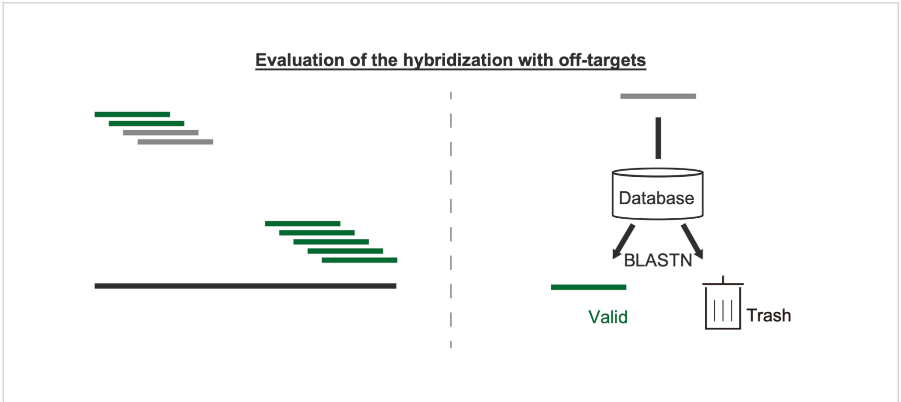


Figure SI6

##### Perform several processes to generate final probes

###### Select Non-overlapping Probe Regions

After specificity filtering, many valid probe regions may overlap with each other (Figure SI7). To maximize probe coverage across the transcript, `hcrkit` selects non-overlapping regions using a greedy algorithm:

1. Sort all valid probe regions by start position (5′ to 3′)
2. Select the first region
3. Skip any regions that overlap with the selected region
4. Select the next non-overlapping region
5. Repeat until all regions are processed

This approach ensures that selected probe regions do not overlap, maximizing the number of probes that can be used simultaneously.

###### Generate Probe Sets

For each selected probe region, `hcrkit` generates a split probe set (P1 and P2) (Figure SI7).

###### Reverse complementation

Because probe regions are extracted from the sense strand, `hcrkit` converts them to antisense sequences (which will bind to the target mRNA) by reverse complementation (Figure SI7).

###### Add initiator sequences

The initiator sequence is split at the specified position (default: 9 nt):

- P1 probe: `[initiator (1–9)] + aa + [reverse_complement(P1 region)]`
- P2 probe: `[reverse_complement(P2 region)] + aa + [initiator (10–21)]`

The "aa" spacer sequences are added at the junction between initiator and binding region (Figure SI1, SI7).

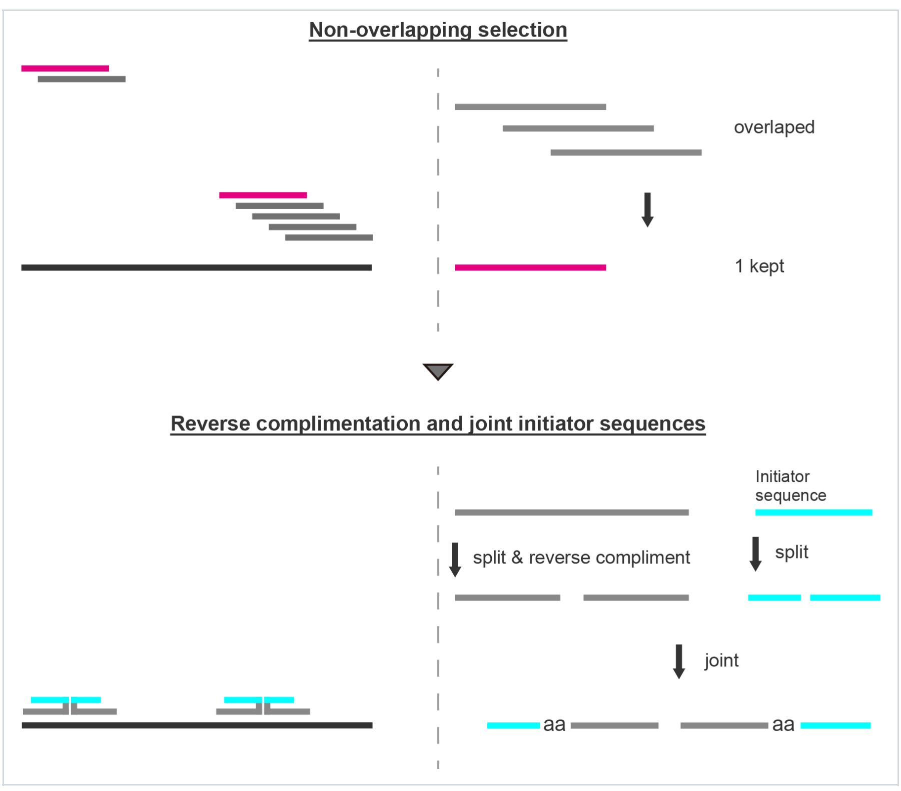


Figure SI7

##### Write Outputs

The final probe sequences are formatted as:

- CSV file: for ordering oligonucleotides
- Summary file: with probe set names and off-target coverage information

This workflow is implemented across `hcrkit.py` (main pipeline) and `core.py` (core functions), working together to automate the entire probe design process.
